# Analysis of Spleen Histopathology, Splenocyte composition and Hematological Parameters in Mice Infected with *Plasmodium berghei* K173

**DOI:** 10.1101/2021.01.22.427878

**Authors:** Huajing Wang, Shuo Li, Zhao Cui, Tingting Qin, Hang Shi, Ji Ma, Lanfang Li, Guihua Yu, Tingliang Jiang, Canghai Li

## Abstract

Malaria is a fatal disease that presents clinically as a continuum of symptoms and severity, which are determined by complex host-parasite interactions. Clearance of infection is believed to be accomplished by the spleen and mononuclear phagocytic system (MPS), both in the presence and absence of artemisinin treatment. The spleen filters infected RBCs from circulation through immune-mediated recognition of the infected RBCs followed by phagocytosis. Using different strains of mice infected with *P. berghei* K173 (PbK173), the mechanisms leading to splenomegaly, histopathology, splenocyte activation and proliferation, and their relationship to control of parasitemia and host mortality were examined. Survival time of mice infected with PbK173 varied, although the infection was uniformly lethal. Mice of the C57BL/6 strain were the most resistant, while mice of the strain ICR were the most susceptible. BALB/c and KM mice were intermediate. In the course of PbK173 infection, both strains of mice experienced significant splenomegaly. Parasites were observed in the red pulp at 3 days post infection in all animals. All spleens retained late trophozoite stages as well as a fraction of earlier ring-stage parasites. The percentages of macrophages in infected C57BL/6 and KM mice were higher than uninfected mice on 8 dpi. Spleens of infected ICR and KM mice exhibited structural disorganization and remodeling. Furthermore, parasitemia was significantly higher in KM versus C57BL/6 mice at 8 dpi. The percentages of macrophages in ICR infected mice were lower than uninfected mice, and the parasitemia was higher than other strains. The results presented here demonstrate the rate of splenic mechanical filtration and the splenic macrophages likely contribute to an individual’s total parasite burden. This in turn can influence the pathogenesis of malaria. Finally, different genetic backgrounds of mice have different splenic mechanisms for controlling malaria infection.

---

*Plasmodium falciparum* parasitescause lethal infections worldwide, especially in Africa (1). Reducing this disease burden continues to rely heavily on the availability and proper use of effective antimalarial drugs. Artemisinin and its derivatives are sesquiterpene lactones with potent activity against nearly all blood stages of *P. falciparum*. There is a natural and complex variation in the pathogenesis and clinical presentation of malaria, which is influenced by host age, immunity and genetic background, as well as by environmental conditions and parasite genetics (2,3). Host immunity and genetic factors are estimated to account for one quarter of the total variability in malaria severity (4,5). Host defense mechanisms such as removal of circulating parasites by the spleen and mononuclear phagocytic system (MPS) are thought to play a major role in rapid control of infection (6), in the presence or absence of artemisinin treatment (7).

The function of the spleen is to remove senescent erythrocytes (RBCs) and circulating foreign material such as bacteria or cellular debris (8). The structure of the spleen is complex with 2 overlapping blood circulations—a rapid flow by-pass, called the fast closed circulation, which accommodates roughly 90% of the splenic blood flow (100–300 mL/min in a healthy adult), and a slow open circulation in which the blood is filtered through narrow inter-endothelial slits (9,10). In the slow open microcirculation, RBCs navigate through the cords of the red pulp before returning to the vascular beds by squeezing between endothelial cells in the sinus walls (11–13). Crossing splenic inter-endothelial slits poses the greatest demand on RBC deformability in the body (14) and is believed to result in the retention of less malleable RBCs or in removal of intraerythrocytic bodies. In malaria, the spleen filters infected RBCs from circulation by physical selection as well as immune-mediated recognition and phagocytosis of infected RBCs (15). These processes play a central role in the clearance of circulating malaria parasites (6). The rate of splenic mechanical filtration may be one factor affecting an individual’s total parasite burden and the pathogenesis of malaria. Understanding the role of the spleen in host defense may shed additional light on the variation in human susceptibility to malaria and offer insights into possible mechanisms of malaria pathogenesis.

In the present study, the host defense against blood-stage malaria was examined by using different strains of mice infected with P. *berghei* K173 (PbK173), a rodent-lethal strain of malaria. Parasitemia and survival were measured to monitor the course of infection in C57BL/6, BALB/C, ICR, and KM mice. Since C57BL/6 mice were found to be more resistant to this infection, parameters indicative of a protective host response to infection were also characterized in the four strains mice. These included splenomegaly, histopathology, splenocyte subsets, hematological parameters.

## RESULTS

### Progression of infection with PbK173 in mice

Mice (18~22g) were purchased from Weitonglihua, and were inoculated with 10^7^ P. *berghei* K173 infected erythrocytes in 200 μL of sterile saline buffer i.p. Mice were monitored daily for any signs of malaria, including parasitemia and behavioral changes, compared to uninfected controls.

The combination of different mouse strains and parasites resulted in different disease outcomes following infection. BALB/c, ICR and KM mice developed infection with parasites observed in the blood as early as 1 dpi, whereas C57 mice converted at 2 dpi. Parasitemia was progressive in all groups (Fig. 1A). On 5 dpi, the parasitemia in ICR mice was 58.6%, 90% on the 8 dpi, and all animals died between 9-11 dpi (Fig. 1B). In BALB/c mice, parasitemia approached 50% by 8 dpi, and peaked at 80% on 10 dpi. The BALB/c mice all succumbed between 16-24 dpi. On the 15th day, the highest parasitemia of the KM mice was 65%, and the animals all died between 15-23 dpi. The parasitemia of C57 mice was close to 50% on the 17th day, and the highest parasitemia reached 80% on the 20th day. All C57 mice died from malaria by 29 dpi.

**FIG 1:**
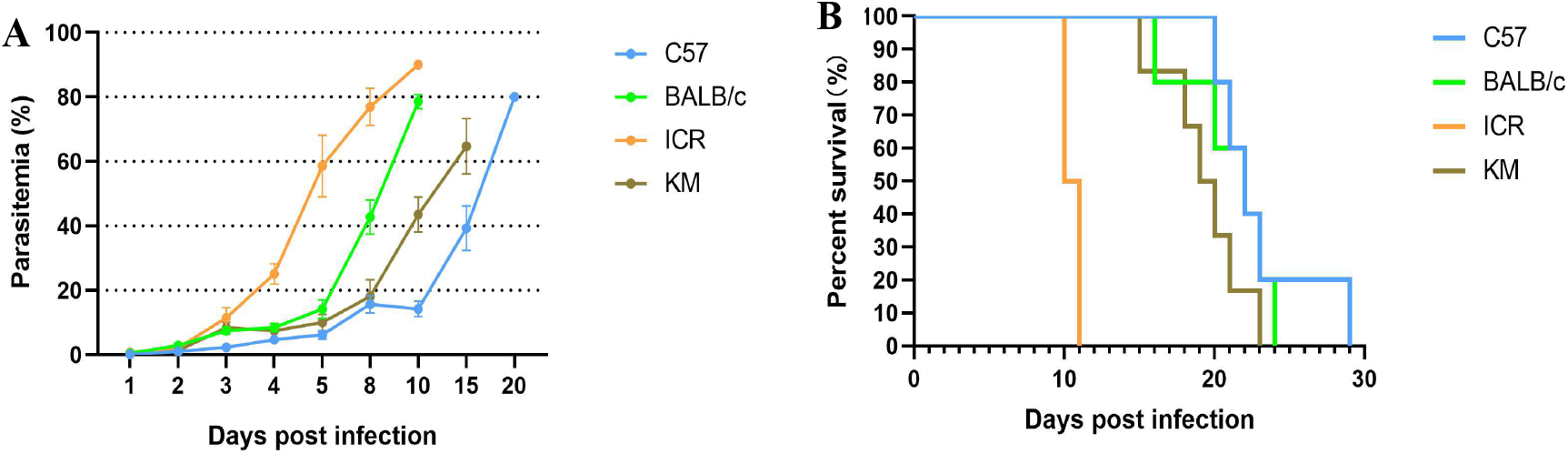
Parasitemia and survival curves in C57BL/6, BALB/c, ICR and KM mice following PbK173 infection. A: The percentage of parasitemia is presented as the arithmetic mean of each mouse strain ± SD. B: Survival of mice without treatment (n=6) as a function of days post infection with PbK173.

It was observed that C57 mice were more resistant to infection than the other strains examined, as evidenced by the latest peak parasitemia and prolonged survival. The highest parasitemia of KM mice was the lowest observed, but its survival was shorter than that of C57BL/6 mice. The highest parasitemia of ICR mice was higher than other strains, although these mice succumbed the fastest.

### Hematological Parameters

In this study, RBC counts (mean ± SD, 10^12^/L) for uninfected control mice were as follows: C57BL/6 (10.01±0.25), BALB/c (7.93±1.01), ICR (8.16±0.16), KM (8.64±0.29). These values are within normal ranges, as reported previously. By 5 dpi,, all strains of mice presented with anaemia, thrombocytopaenia, and leukocytosis (Table 1).

**Table 1:** Effects of PbK173 infection on some hematological parameters in infected mice on 5 dpi.

| Hematological parameters | C57BL/6 |  | BALB/c |  | ICR |  | KM |  |
| --- | --- | --- | --- | --- | --- | --- | --- | --- |
|  | Uninfected | Infected | uninfected | infected | uninfected | infected | uninfected | infected |
| PLT (10<sup>9</sup>/L) | 1278.83±89.60 | 285.00±34.88↓*** | 1310.50±57.76 | 312.33±19.59↓*** | 1635.67±68.14 | 296.00±33.80↓*** | 1476.17±99.99 | 341.33±31.87↓*** |
| RBC (10<sup>12</sup>/L) | 10.01±0.25 | 7.75±0.62↓*** | 7.93±1.01 | 6.62±0.37↓* | 8.16±0.16 | 3.39±0.35↓*** | 8.64±0.29 | 4.44±0.34↓*** |
| HGB (g/L) | 149.67±2.66 | 113.83±8.45↓*** | 132.33±12.07 | 101.50±4.42↓* | 137.17±4.26 | 60.00±6.96↓*** | 129.67±5.98 | 68.83±4.92↓*** |
| HCT (%) | 47.23±0.96 | 37.78±3.03↓*** | 40.40±3.28 | 32.58±1.33↓* | 42.48±1.10 | 21.27±2.19↓*** | 42.67±2.02 | 24.77±1.86↓*** |
| MCV (fL) | 46.52±0.43 | 48.40±0.71↑*** | 45.02±0.67 | 49.22±1.48↑* | 51.77±1.33 | 62.72±3.15↑*** | 48.75±0.83 | 56.26±1.12↑*** |
| MCH (pg) | 14.71±0.07 | 14.56±0.24 | 14.73±0.15 | 15.46±0.63↑* | 16.86±0.35 | 17.55±0.76 | 15.18±0.35 | 15.75±0.12↑* |
| MCHC (g/L) | 316.83±2.04 | 301.17±2.14↓*** | 327.17±4.35 | 310.50±5.17↓*** | 321.67±3.26 | 283.83±5.49↓*** | 310.67±3.38 | 279.00±5.66↓*** |
| RDW-CV (%) | 17.35±.26 | 16.20±0.83↓* | 19.03±0.99 | 16.72±0.92↓* | 15.85±0.32 | 19.97±1.21↓* | 16.75±0.57 | 15.83±1.64 |
| RDW-SD (fL) | 22.80±0.59 | 24.50±1.16↑* | 25.82±2.22 | 26.87±1.49 | 26.70±0.84 | 41.11±4.13↑*** | 27.27±0.74 | 31.98±1.85↑*** |
| WBC (10<sup>9</sup>/L) | 2.74±0.39 | 5.84±2.16↑* | 4.37±0.93 | 8.64±1.95↑* | 2.52±0.35 | 5.63±1.64↑* | 1.65±0.30 | 3.45±0.43↑*** |
| NEUT (%) | 10.13±0.4 | 88.33±5.83↑*** | 32.68±9.50 | 76.53±10.66↑*** | 16.50±1.07 | 91.65±2.59↑*** | 14.78±1.31 | 94.77±1.04↑*** |
| LYMPH (%) | 82.80±2.78 | 5.71±3.13↓*** | 52.68±9.16 | ---↓*** | 73.98±1.77 | ---↓*** | 74.57±2.09 | ---↓*** |
| MONO (%) | 6.35±1.55 | 5.68±2.45 | 2.10±0.38 | 7.00±2.46↑* | 4.66±0.85 | 2.71±0.85↓* | 4.85±1.01 | 3.11±0.46↓* |
| EO (%) | 1.47±0.67 | 0.47±0.38↓* | 2.13±0.59 | 1.33±0.68 | 4.06±0.53 | 3.98±2.73 | 4.60±1.66 | 0.71±0.24↓* |
| BASO (%) | 0.32±0.23 | 0.63±0.23 | 0.16±0.05 | 0.38±0.04↑* | 0.25±0.20 | 0.45±0.08 | 0.51±0.17 | 0.58±0.20 |

Values are given as mean ± SD, n=6. \* and \*\*\* indicate $p < 0.05$ and $p < 0.001$ compared with the uninfected group, respectively

Malaria infections induce lymphocytopenia and an increase in neutrophils, which is indicative of systemic inflammation (16). In all 4 strains of mice, the percentages of lymphocytes decreased compared to the baseline values starting from 1 dpi. In ICR mice, the decrease progressed significantly over the first 3 days. The lymphocytopenia progressed slowly in BALB/c and KM mice on the 2nd-3rd day of infection (Fig. 2A). In all mice, the percentages of neutrophils increased relative to uninfected control mice starting from 1 dpi (Fig. 2B).

**FIG 2:**
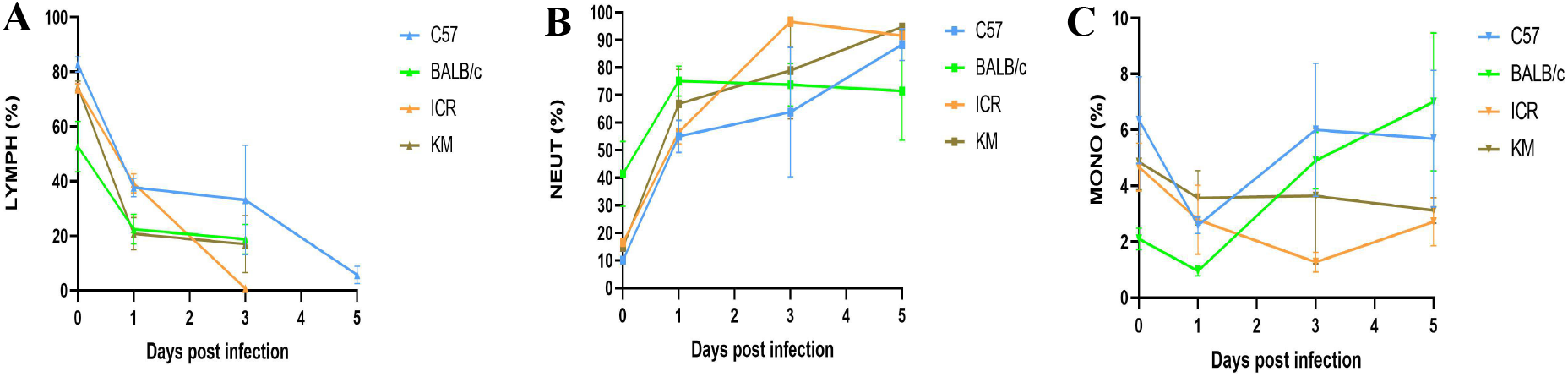
Effects of PbK173 infection on hematological parameters in C57BL/6, BALB/c, ICR and KM mice. A: Lymphocytes B: Neutrophils C: Monocytes.

**FIG 3:**
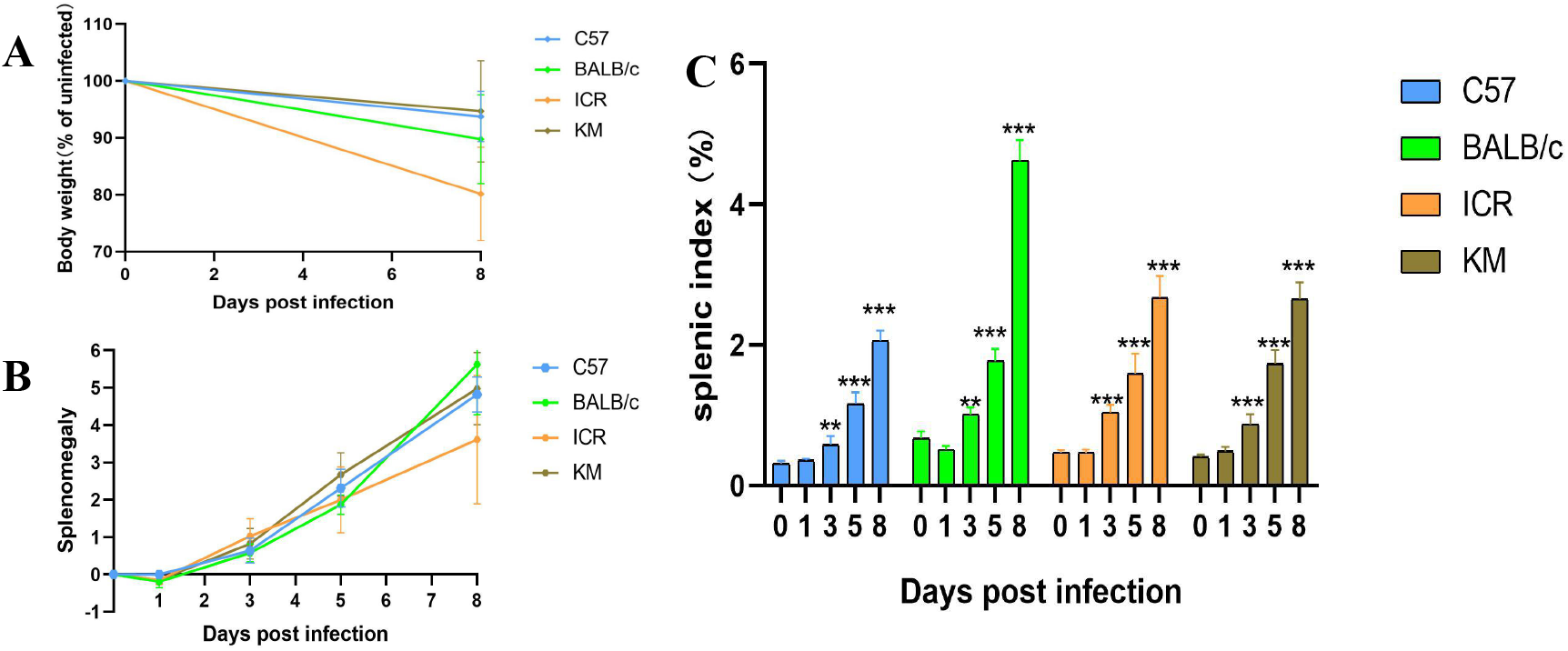
The change in body weight (A) and spleen weight (B) in the four strains of mice, spleen weights were normalized to the weight of uninfected groups (%). C: Splenic index in uninfected and PbK173 infected mice. Spleen weights of infected mice were determined on days 3, 5, and 8 post infection. The splenic index was determined as the ratio of spleen weight to body weight. All data are presented as mean ± SD, n=6. ** and *** indicate statistical significance at *p* < 0.01 and *p* < 0.001 compared with the uninfected group, respectively.

The percentages of monocytes decreased compared with baseline values on at 1 dpi for all infected groups. But on 5 dpi, the percentages of monocytes in BALB/c (7.00±2.46, *p* < 0.05) and C57BL/6 (5.68±2.45, *p*>0.05) mice were increased.

In contrast, monocyte counts decreased in ICR (2.71±0.85, *p* < 0.05) and KM (3.11±0.46, *p* < 0.05) mice (Fig. 2C).

### Gross and histopathologic analysis of the spleen

The spleen is an important site of erythropoiesis, the clearance of infected RBCs (iRBCs), and immune system activation in response to blood-stage malaria (17). In the present study, the body weight of infected mice was observed to decline following infection with PbK173. The spleen weight of infected groups was observed to increase beginning at 3 dpi. In response to infection, all mice experienced significant splenomegaly, but the splenic index was significantly higher in infected BALB/c mice.

Figure 4 shows the histopathological sections of the spleen tissue. A clear distinction between the red and white pulp, resting follicles, and marginal zones were evident in the spleen of normal uninfected control mice (Fig. 4: A, E, I, M). Severe congestion and enlarged red pulp was observed in spleens of infected mice at 3 dpi (Fig. 4: B, F, J, N). To a similar all mouse strains examined, increases in red and white pulp cellularity was observed, and the clear marginal zones surrounding follicles became inapparent (except in C57BL/6) at 8 dpi (Fig. 4: C, G, K, O). Furthermore, extensive vacuolation in the red pulp at 8 dpi was observed in spleens from ICR and KM mice (Fig. 4: L, P).

**FIG 4:**
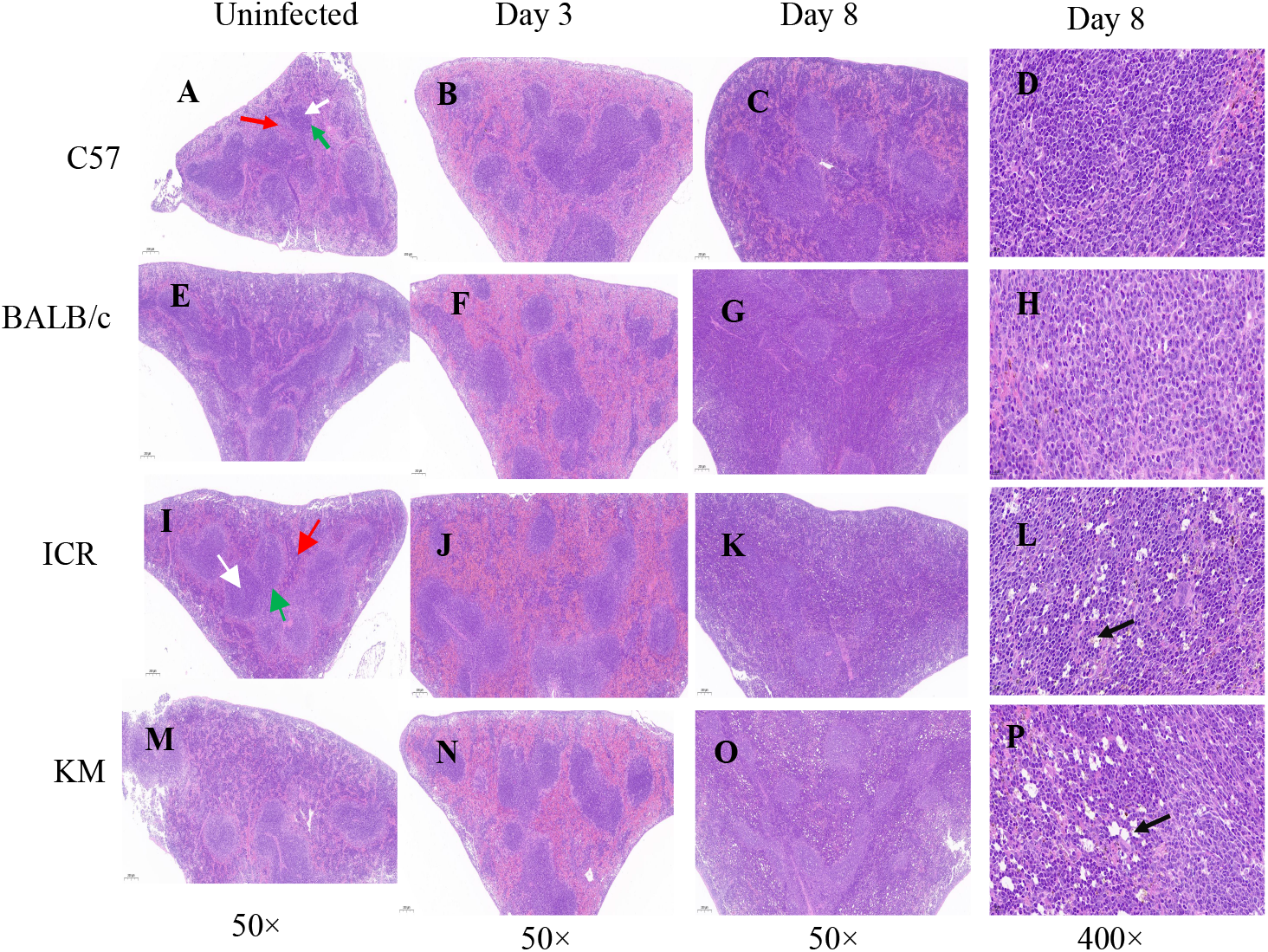
H&E stained sections of spleen from infected or uninfected mice. white arrows: white pulp; red arrows: red pulp; green arrows: marginal zone; black arrows highlight vacuolation.

Spleen sections were prepared and examined via bright-field microscopy coupled with hematoxylin and eosin (H&E) staining (Fig. 5). Parasite pigments in the pulp histiocytes and sinusoidal lining cells were observed at 3 dpi in all PbK173 infected mice. The extent of malaria pigmentation in the spleen is correlated with high parasitemia. The spleens of C57BL/6, BALB/c, ICR, and KM mice filtered late trophozoite stages as well as a fraction of earlier ring-stage parasites out of the blood at 3 dpi (Fig. 5: B, E, H, K). However, KM mice only retained the late trophozoite stage on day 8 (Fig. 5I). This phenomenon may be due to changes of the splenic structure, which could have resulted in alterations to the filtering function of the spleen. Transformations in the red pulp and splenic vasculature may modulate the mechanical retention threshold and regulate the microcirculatory trapping of blood cells in the spleen.

**FIG 5:**
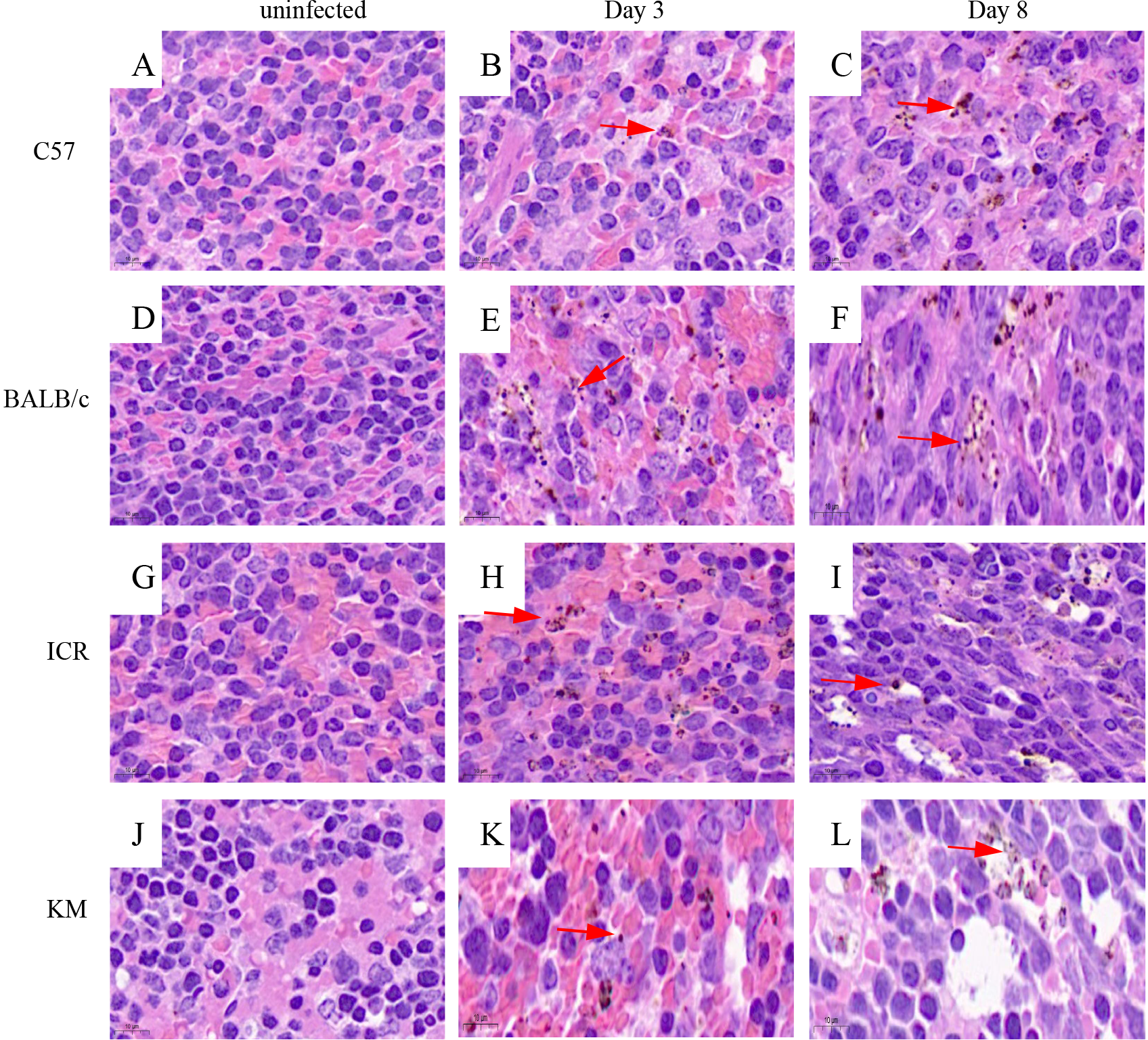
H&E staining of spleen from control and PbK173 infected mice (1000×). Red arrows indicate malarial pigments, which appear as small brown punctate staining.

### Analysis of Splenocyte Subsets

Next, the distribution of macrophage and T lymphocyte subpopulations were analyzed. Using single-cell suspensions from PbK173 infected or uninfected control spleens, flow cytometry was performed to quantify total leukocytes (CD45^+^ cells), total T lymphocytes (CD45^+^CD3^+^ cells), T cell subsets (CD4^+^ and CD8^+^ cells), and monocytes/macrophages (CD45^+^F4/80^+^ cells) (Fig. 6). A significant decrease in total leukocyte in both C57BL/6, BALB/c, ICR and KM infected mice (*p*<0.01) was observed (Fig. 6A). The data also indicated a significant decrease in the percentages of the T lymphocytes (CD3^+^ cells, except ICR and KM infected mice) (Fig. 6B) and CD8^+^ cells (Fig. 6D), while the percentages of CD4^+^ cells (Fig. 6C) in both C57BL/6, BALB/c, ICR and KM infected mice increased.

**FIG 6:**
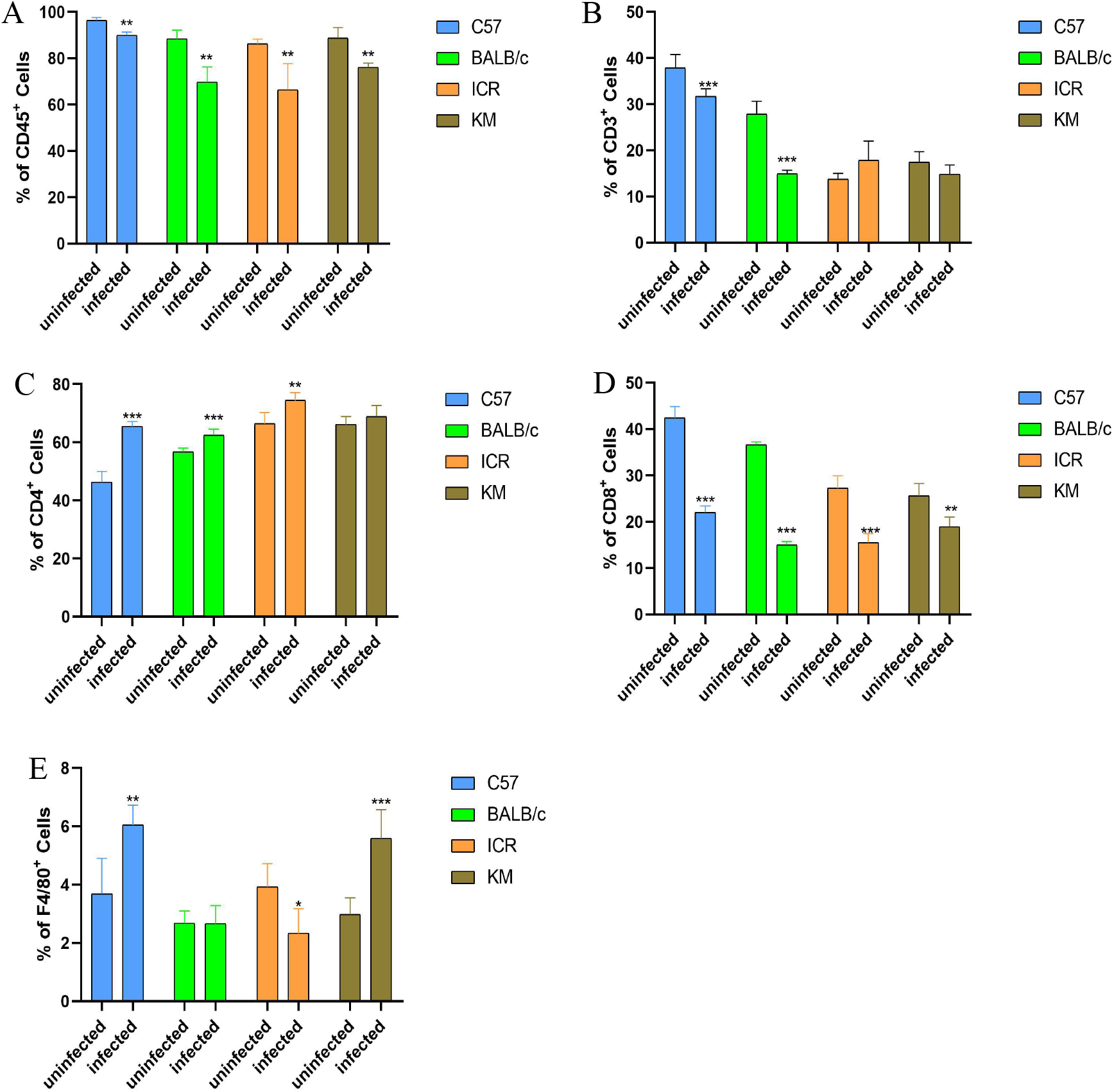
Flow cytometry analysis of splenocyte subsets at 8 dpi. Splenocytes of mice infected with parasites were incubated with the required antibodies according to the manufacture’s protocol for antibody dilution, incubation duration, etc., and were analyzed in a CytoFLEX flow cytometer. *, ** and *** indicate statistical significance at *p* < 0.05, *p* < 0.01 and *p* < 0.001 compared with the uninfected group, respectively.

The trends in macrophage percentages differed between the 4 strains of mice. The percentage of macrophages did not change significantly in BALB/c infected mice. A decrease in the percentage of macrophages was however observed in the infected ICR mice. Conversely, a significant increase in the percentages of the macrophages in C57BL/6 and KM infected mice as compared to uninfected mice was observed (Fig. 6E).

Splenic red pulp macrophages, located between the splenic cords and venous sinuses, are well positioned to clear iRBCs and are important for controlling blood-stage malaria (18). In the present study, after infection with PbK173, the percentages of macrophages in the spleen of C57BL/6 mice exhibited the greatest increase of all the mouse strins (Fig. 7B), and the parasitemia progressed the slowest compared to the other mouse strains (Fig. 7A).

**FIG 7:**
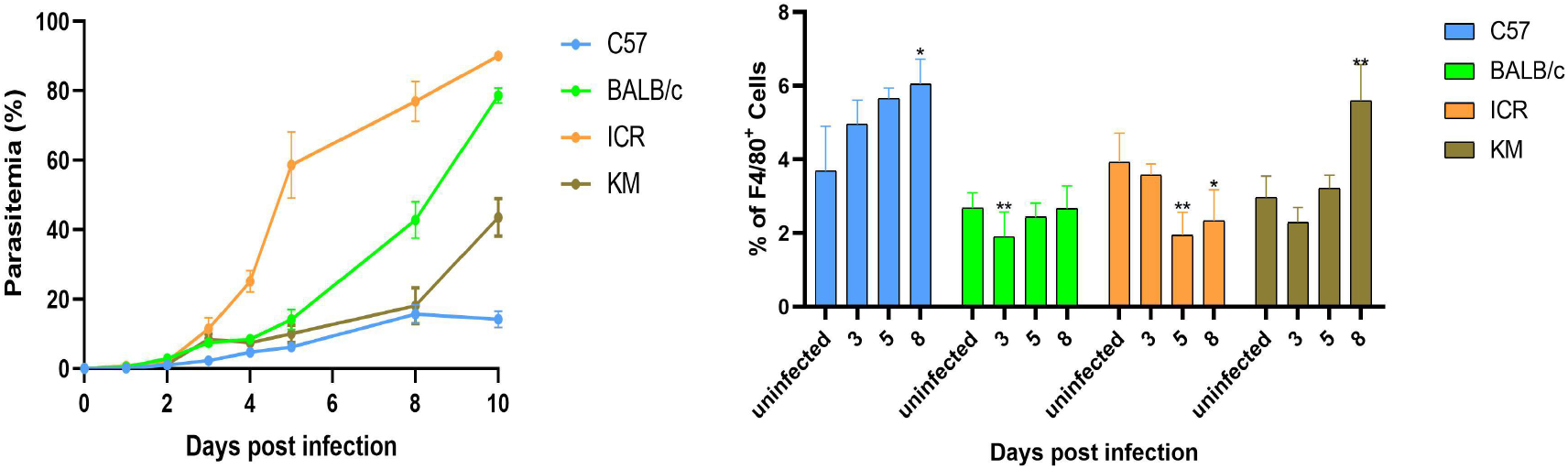
Parasitemia and flow cytometry analysis of macrophages in C57BL/6, BALB/c, ICR and KM mice over the course of PbK173 infection. * and ** indicate *p* < 0.05, *p* < 0.01 compared with the uninfected group, respectively.

The percentage of macrophages in the spleen of KM mice was higher than uninfected group on 8 dpi. The spleens in KM mice exhibited structural disorganization and remodeling, which likely affected the mechanical retention threshold. As a result, only late trophozoite stages were retained, which led to a significantly higher parasitemia in KM than C57BL/6 mice. The percentages of macrophages in ICR infected mice were lower than uninfected mice, and the parasitemia was higher than other strains during the course of PbK173 infection (Fig. 7).

These data show that the rate of splenic mechanical filtration and the splenic macrophages may be important factors in determining an individual’s total parasite burden and potentially influencing the pathogenesis of malaria, and different genetic backgrounds of mice have different mechanisms for controlling malaria infection in the spleen.

## DISCUSSION

In this study, the survival time of mice infected with PbK173 varied, although the infection was uniformly lethal. Mice of the C57BL/6 strain were the most resistant, while ICR mice were the most susceptible. Both BALB/c and KM mice were intermediate.

Changes in blood cell counts are characteristic of plasmodium infection. Haematological changes during the course of a malaria infection, such as anaemia, thrombocytopaenia, leukocytosis, and leucopenia are defining. Malaria infections also induce lymphocytopenia, which is accompanied by an increase in neutrophil count, and is a sign of systemic inflammation. Neutrophils (also known as polymorphonuclear cells) are the most common white blood cell in the body (19) and are one of the immune system’s first lines of defense against infection. They can neutralize pathogens through several mechanisms, including phagocytosis, the production of reactive oxygen species (ROS) and other antimicrobial products, or by the formation of neutrophil extracellular traps (NETs) (20). Additionally, they also play a role in the activation and regulation of the immune response through the secretion of cytokines and chemokines (21), and the presentation of foreign antigens (22). A substantial body of work has been conducted to investigate the role of neutrophils in malaria. It has been determined that during infection, neutrophils are activated and are capable of clearing malaria parasites by a number of mechanisms.

In this experiment, the above hematological parameters of different mouse strains exhibited similar trends with respect to changes in the parameters, although the changes in the percentages of monocytes varied. Cells of the monocyte/macrophage lineage are one of the main sources of cytokines in malaria-infected individuals. While some of the cytokines may be of importance for parasite clearance (eg, IL-12) (23), others may be major contributors to disease progression (eg, TNF) (24). Monocytes recognize *Plasmodium falciparum* biological products and Pf-infected erythrocytes directly through pattern recognition receptors (PRR) (25), as well as complement- or IgG-opsonized erythrocytes and parasite components via complement receptors and Fcγ receptors (26). Activated monocytes have several important effector functions in the host defense against malaria, including phagocytosis (27), cytokine production (28), and modulation of adaptive immune responses (29,30). At 5 dpi, compared with the uninfected groups, the percentages of monocytes in BALB/c and C57BL/6 mice were elevated, whereas the values in ICR and KM mice were reduced. Conversely, the parasitemia in ICR mice was significantly higher than KM mice. These results indicate that perhaps the immune cells in peripheral blood are only part of the host response necessary to control plasmodium infection.

The spleen is a key site for removal of parasitized red blood cells, generation of immunity and production of new red blood cells during malaria. The importance of the spleen for the control of malaria was confirmed by studying the response of splenectomized humans and rodents to infection. Humans with acute *P. falciparum* malaria who had previously undergone splenectomy had decreased clearance of iRBCs from the circulation (31). The mice subjected to partial splenectomy presented a level of parasites similar to that of non-splenectomized mice, while the animals subjected to full splenectomy had twice the amount of circulating parasites (32). Furthermore, parasite clearance after drug treatment was delayed in splenectomized patients, with RBCs containing dead parasites being retained in the circulation for prolonged periods, compared with individuals with a functional spleen (33).

During acute attacks of malaria, splenomegaly is one of the typical signs of malaria, and the degree of splenomegaly often impacts the host’s ability to mount a successful response to the parasite (34). Besides an increase in the organ volume and mass, the spleen also exhibits structural disorganization and remodeling. These changes include expansion of the red pulp, transient loss of the marginal zone, increased vasculature, and activation of barrier cells, which may establish a blood-spleen barrier that can drastically alter splenic blood circulation (35,13,36). In this study, the spleen index of infected groups was observed to increase from 3 dpi. Severe congestion and enlarged red pulp was evident in the infected mice. By 8 dpi, infection-induced increases in red and white pulp cellularity and the marginal zones surrounding follicles disappeared in all strains of mice examined. However, the spleens of C57BL/6 and BALB/C infected mice maintain their structural integrity integrity, although the spleen index of BALB/C changed the most. The spleen of ICR and KM mice exhibited severe vacuolation, and the splenic structure was highly atypical, with many of the features absent at this time. This could be a result of the spleen structures of mice with different genetic backgrounds possessing different tolerances and pathologies to infection with malaria.

During the erythrocytic stages of malaria infection, the spleen plays a critical role in the host immune response. Elimination of infected erythrocytes occurs through activation of cellular and humoral immune responses, and through mechanical filtration. White and red pulp structures have specific functions in the human spleen. The white pulp is a major control center for the humoral immune response, especially to circulating antigens. The red pulp exerts a unique and subtle control of the surface integrity and biomechanical properties of erythrocytes. To be left in circulation, RBCs must be fit enough to cross a very specific structure of red pulp sinuses, the interendothelial slit (IES). Older erythrocytes, or those modified by innate or acquired conditions, are eventually retained in the splenic red pulp and processed by red pulp macrophages (RPMs) (37).

During asexual replication (including the sequential ring, trophozoite, and schizont stages), parasite maturation induces changes in the host RBC with novel proteins synthesis (38,39). As the parasite develops, the infected RBC (iRBC) loses its biconcave shape and progressively becomes spherical and rigid (40). Furthermore, the surface area-to-volume ratio decreases, the shear elastic modulus of the plasma membrane, and the cellular viscosity increase (41). The loss of RBC deformability is not limited to mature stages, but starts soon after parasite invasion. During the ring stage (within the first 16–24 h after RBC invasion), iRBC undergo up to 9.6% surface area loss (42,43). The altered deformability of the plasmodium-infected RBC may result in increased retention in the spleen. More than 50% of ring-iRBC are retained upon *ex vivo* transfusion through human spleens (43). These retention and accumulation processes stem from the splenic screening of RBC deformability (44). However, no direct evidence exists demonstrating the correlation among the rate of splenic mechanical filtration, macrophages, and infection severity. In this study, at 3 days post PbK173 infection, malaria pigments were observed in the red pulp in great abundance. The pigments consisted of parasites in the ring and trophozoite stages.

In a systemic pathological study of cerebral malaria in African children, enlarged spleens and abundant malaria pigments in splenic macrophages were observed in the majority of the 103 fatal cases. These observations point to an important role of the spleen in parasite control. In this study, the number of macrophages in the spleen of C57BL/6 mice infected with PbK173 was higher than that in uninfected controls, and the parasitemia was lower than other strains during. The percentage of macrophages in the spleen of infected ICR mice was lower than the uninfected group, and the parasitemia increased the fastest. During the infection period, the ratio of macrophages in the spleen of BALB/c mice was not significantly different from that of the uninfected group. The growth rate of the parasitemia was lower than in ICR mice, but higher than that of C57BL/6 mice. This can be explained by the fact that macrophages complement the filtering function of the spleen to control parasitic infections.

Artemisinin-based combination therapies (ACTs) are the standard of care to treat uncomplicated falciparum malaria. However, resistance to artemisinins was first reported through observations of a 100-fold reduction in parasite clearance rate in Pailin, Western Cambodia, in 2009 (45). The clearance rate is defined as a parasite clearance half-life ≥5 hours following treatment with artesunate monotherapy or an ACT (46). Unfortunately, this has become a common issue in Southeast Asia (SEA). Artemisinin resistance has been associated in with multiple nonsynonymous single nucleotide polymorphisms (NS-SNPs) in the propeller domain of the gene encoding the *P. falciparum* K13 protein (K13PD) (47,48). In Africa, significantly prolonged clearance has yet to be observed (49).

The delayed clearance phenotype initially observed in SEA does not represent “drug resistance” in the traditional use of the term. Antimicrobial drug resistance is typically defined by quantifiable shifts in the cytostatic or cytocidal potency of the antimicrobial drug. It is important to emphasize that as a 3-day course of artemisinin has never been considered a curative regimen. Artemisinins remain effective, even if they require a longer treatment course or other modifications to the combination-treatment regimen (50). It is not clear if a delay in parasite clearance with artemisinin treatments be defined as drug “resistance.”

The spleen controls malaria infection by removing the plasmodium from the blood. Splenic functions of mice with different genetic backgrounds vary, and when splenomegaly occurs after malaria infection, the prognosis is not favorable. However, it is not clear if splenomegaly affects the organ’s ability to clear parasites from the blood.

In summary, the filtering function of the spleen and the expression of macrophages may also be involved in the control of malaria infection. Mice with different genetic backgrounds have different filtering functions in the spleen from the expression of macrophages. The tolerance is also different, which may be the reason for the differential susceptibility of different strains of mice to PbK173.

## MATERIALS AND METHODS

### Parasite strains and culturing conditions

*Plasmodium berghei* K173, a gift from Dr. Dai of Chengdu University of TCM, was serially passaged *in vivo* in mice. Infected blood was harvested at day 5–7 post-infection and stored as frozen stabilates in Alsever’s solution containing 10% glycerol.

### Mice and infection

Male C57BL/6, BALB/C, ICR, and KM wild-type (WT) mice (18~22g,) were used in this study. Animals were purchased from Weitonglihua (Beijing, China). A total of 6 mice per group were infected intraperitoneally with 10^7^ PbK173-infected RBCs and were provided water and standard laboratory mouse chow diet *ad libitum* throughout the experiment. All mice were housed in pathogen-free animal facilities at the Institute of Chinese Materia Medica, China Academy of Chinese Medical Sciences.

### Measurement of hematologic parameters and parasitemia

Complete blood counts were obtained with a XN-1000V [B_1_] blood analyzer (Sysmex, Japan). Parasitemia (i.e., the percentage of infected RBCs) was assessed at each time point by microscopic counts of thin blood smears stained with Giemsa solution (Sigma -Aldrich, USA).

### Isolation of immune cells from mouse spleen

Spleen samples were surgically removed and weighed in a sterile hood. One part of each spleen sample was removed and fixed 4% paraformaldehyde for histopathologic examination, and the remainder was used for isolation of splenocytes.

Spleens harvested under aseptic conditions were ground into small pieces and passed through a sterilized 200 mesh to prepare crude splenocyte suspensions at room temperature. Samples were then centrifuged at 1000 rpm for 8 min at 4 °C, and the remaining splenocyte suspension was re-suspended in red blood cell Lysis Buffer (Thermo Fisher Scientific, USA). After a 10 min treatment, 1× PBS was added to dilute the samples, and then centrifuged at 1000 rpm for 8 min at 4 °C. The pelleted splenocytes in each group were washed twice and adjusted to concentrations of 5 × 10^6^ cells/mL with 1× PBS.

### Analysis of Splenocyte Subsets

The single cell splenocyte suspensions were stained with the following anti-mouse antibodies: CD45-KO525, CD3-FITC, CD4-PC5.5, CD8-APC, and F4/80-PE (Proteintech Group, USA). Splenocytes were incubated with monoclonal antibodies in the dark for 30 min at 4 ℃. According to the manufacturer’s instructions, the specificity of labeling was confirmed by isotype-matched antibody staining controls. The labeled cells were analyzed using a CytoFLEX flow cytometer (Beckman coulter, USA).

### Histological examination

Spleen tissues were fixed in 4% paraformaldehyde, dehydrated through graded alcohol, embedded in paraffin, sectioned at a thickness of 3 μm, and then stained with hematoxylin & eosin (H&E) and Giemsa solution according to standard procedures. Then, the stained slides were mounted in neutral balsam and covered with coverslips. Histopathologic changes were observed by light microscopy (BX43F Olympus, Japan).

### Statistical Analysis

Data were analyzed using SPSS 19.0 (IBM,USA) and reported as mean ± SD. Significant differences between groups were analyzed using one-way ANOVA, and are designated as follows: * *p* < 0.05, ** *p* < 0.01 and *** *p* < 0.001 relative to the uninfected control groups. Survival curves were calculated using GraphPad Prism 8.0 (GraphPad Software,USA).

## ETHIC STATEMENT

Experimental protocols were approved by the Laboratory Animal Ethics Committee of the Institute of Chinese Materia Medica, China Academy of Chinese Medical Sciences (license number SCXK 2016-0006).

## ACKNOWLEDGMENTS

This research was funded by the National Natural Science Foundation of China (81641002 and 81841001). We gratefully acknowledge the help from researchers at the Tang Center for Herbal Medicine Research for providing experiment guidance. We would also like to thank Dr. Dai from Chengdu University of TCM for providing the *Plasmodium berghei* K173 and providing technical guidance.

